# Effect of environmental conditions on seed germination and seedling growth in *Cuscuta campestris*

**DOI:** 10.1101/2024.06.14.598959

**Authors:** Koki Nagao, Taku Takahashi, Ryusuke Yokoyama

## Abstract

The dodder, *Cuscuta*, is an obligate parasitic plant that cannot survive without a host plant and causes major damage to crop yields. To know its growth characteristics before parasitism, we examined the effect of various environmental conditions on seed germination and seedling growth in *Cuscuta campestris*. As for the effect of light on germination, far-red light was rather preferable to red light and the reversible response of the seeds to red and far-red light was confirmed, implicating a phytochrome-mediated signaling opposite to that in many seed plants. Among amino acids, aspartic acid and alanine had a promotive effect and histidine had an inhibitory effect on germination. We further found that, in addition to gibberellic acid, methyl jasmonate was stimulatory to germination and shoot elongation. While 2,4-D extended the viability of trichomes around the root cap, kinetin induced the formation of scale leaves on the shoot and calli at the base of the shoot and the root tip. RT-PCR experiments confirmed that expression of a putative *RbcS* gene for photosynthesis showed no response to light but that of a *Phytochrome A* homolog was increased in the dark. Our results indicate that some of the molecular mechanisms to respond to light and hormone signals are uniquely modified in dodder seedlings, providing a clue for understanding the survival strategy of parasitic plants.

## Introduction

*Cuscuta* species (dodders) are among the most widespread parasitic weeds that attach to and feed on the stem of various types of herbaceous angiosperms. Understanding the molecular basis of parasitism by dodders is critical to develop effective pest control systems since they heavily damage many important crops. There are increasing number of studies about the development of the haustorium, a root-like structure that penetrates the host’s tissue and draws water and nutrients (Yoshida et al. 2016; Shimizu and Aoki 2019; Jhu and Sinha 2022; Hartenstein et al. 2023). Elongation of the haustorium is promoted by host-produced ethylene (Narukawa et al. 2021). Involvement of ethylene in haustorium formation has also been shown in a facultative root parasite in the Orobanchaceae family, *Phtheirospermum japonicum* (Cui et al. 2020). While blue and far-red light strongly enhances the process of parasitism, as-yet-unidentified host-derived signaling factors may be important for the establishment of parasitic connection between dodders and host plants. Transcriptome analysis in *Cuscuta pentagona* revealed that expression of the genes for phytohormone transporters and those involved in cell wall modifications are induced in the process of parasitism (Ranjan et al. 2014). A recent study reported that, in field dodder, *C. campestris*, the genes for cell wall degradation show different expression levels in response to different polysaccharide composition of the host cell wall (Bawin et al. 2024). According to the small-RNA sequencing experiments, some dodders’ microRNAs could target the host genes and further improve the parasitism (Shahid et al. 2018). After establishment of parasitism, some dodder species may synchronize their flowering with that of their hosts. *C. australis* does not have an autonomous flowering behavior but utilizes host-produced florigen as an interplant mobile signal for synchronous floral induction (Shen et al. 2000). On the one hand of the importance of host-derived signals in enhancing the parasitism, an axenic in vitro culture system that enables the entire lifecycle has been developed in *C. campestris* (Bernal-Galeano et al. 2022).

In contrast to the host-dependent growth during parasitism, the growth before parasitism depends mainly on the starch stored in the seedlings as an energy source. However, the developmental control and physiological traits of dodder seedlings in the autotrophic growth phase remain rather unexplored. They emerge a thread-shaped hypocotyl with no cotyledons, which show nastic movements in search of potential hosts. Most dodders form only rudimentary roots with root apices surrounded by a circle of root hair-like trichomes. The shoots have an apical meristem but lack mechanical tissues (Toma et al. 2005; Sherman et al. 2008). Xylem bundles show scattered or circular arrangement, depending on species (Toma et al. 2005). A study of characterization of plastids in six *Cuscuta* species indicated that they show functional diversity, ranging from intact chloroplasts to plastids with impaired protein production and gene expression or reduced plastome gene content (van der Kooij et al. 2000). Comparative analysis of plastid genomes revealed that *Cuscuta* species lost the photosynthesis ability to various extents because of the gradual loss of genes involved in photosynthesis (Wang et al. 2023). Most dodder seedlings become senescent by 7 to 10 days and collapse completely by 2 weeks post-germination. It is thus essential for their survival to find out host plants in this short autotrophic phase.

To know developmental and physiological characteristics of dodders prior to parasitism, the current study focuses on the response of *C. campestris* seeds and seedlings to a variety of external conditions. Our results provide a basis for further understanding the survival strategy of dodders and the pest control against them.

## Materials and Methods

### Plant Materials and Growth Conditions

Seeds of *C. campestris* (Narukawa et al. 2021) were harvested from the plants parasitizing *Arabidopsis thaliana*, the Columbia-0 ecotype, grown under a 16/8 h light/dark cycle at 25 °C. Dry *C. campestris* seeds were treated by soaking in concentrated sulfuric acid for 15 □min, rinsed three times with distilled water, sterilized in bleach solution containing 1% sodium hypochlorite for 10 min, and washed five times with distilled water. The seeds were then sown on 1% agar plates containing 1x Murashige and Skoog (MS) salts and 1% (w/v) sucrose and incubated for 3 days under a 16/8 h light/dark cycle of fluorescent white light (30 µmol m^−2^ s^−1^) at 25 °C, unless otherwise stated. For seedling growth experiments, the germinating seedlings that show full emergence of the root at 3 days after sowing were transferred to new plates and grown for 7 days under each test condition.

For light irradiation experiments, plates were placed under exposure of approximately 7.5 µmol m^−2^ s^−1^ of blue light (λ_max_ = 450 nm, Tokulumi Company Limited, China), red light (λ_max_ = 660 nm, Tokulumi Company Limited), or 5 µmol m^−2^ s^−1^ of far □red light (λ_max_ = 720 nm, Fuji electronic, Japan). For pH experiments, the MS media added with 10 mM final concentration of tris base were modified with 1 N HCl. For temperature treatments, plates were placed face down to reduce the effect of humidity changes on the growth.

### Microscopy

Seedlings were imaged with a stereomicroscope (SMZ1500, Nikon, Japan) and the shoot length was analyzed with the ImageJ software. For tissue sections, samples were fixed overnight in 50% ethanol, 5% formaldehyde and 5% acetic acid, dehydrated through an ethanol series, and embedded in Technovit 7100 resin (Heraeus Kulzer, Germany) according to the manufacturer’s instructions. Sections of 10 µm thickness were stained with 0.1% toluidine blue and observed under a light microscope (ECLIPS 80i, Nikon).

### RNA extraction and qRT-PCR

For Total RNA was prepared from seedlings using a NucleoSpin RNA Plant Kit (Takara, Japan) according to the manufacturer’s instructions and reverse-transcribed using a PrimeScript RT reagent Kit (Takara) with the oligo-dT primer. The resulting first-strand cDNA was used as a template for real-time PCR with gene-specific primers. PCRs were performed using a KAPA SYBR FAST qPCR Kit (KAPA Biosystems, USA) with the Thermal Cycler Dice Real Time System (Takara). *EF1a* was used as an internal standard. Specific primers used for each cDNA amplification are shown in Fig. S1.

### Chemicals

MS salts, sugars, amino acids, and polyamines except thermospermine were purchased from Fujifilm Wako (Japan). Thermospermine was purchased from Santa Cruz Biotechnology (USA). All plant hormones and bikinin were obtained from Tokyo Chemical Industry (Japan).

### Statistical analysis

Three replicates of 20 seeds for germination and 10 seedlings for shoot growth were examined in each treatment. The data were analyzed by Student’s *t-*test between control and experimental groups or oneway ANOVA with Tukey–Kramer multiple comparison test.

## Results

### Far-red light enhances seed germination

After treatment with concentrated sulfuric acid, seeds were sown on MS agar plates. Under our experimental conditions, the germination rate reached close to maximum in 6 days after sowing (Fig. 1A). In this study, the seeds were judged as germinated based on visible root protrusion (Fig. 1B). The germinating seedlings were transferred at 3 days after sowing to new agar plates and grown under a 16/8 h light/dark cycle of fluorescent light. The shoot length reached a maximum level at 10 days after sowing (7 days after transfer; Fig. 1C). A typical morphology of 10-day-old seedlings is shown in Fig. 1D.

**Fig. 1.**
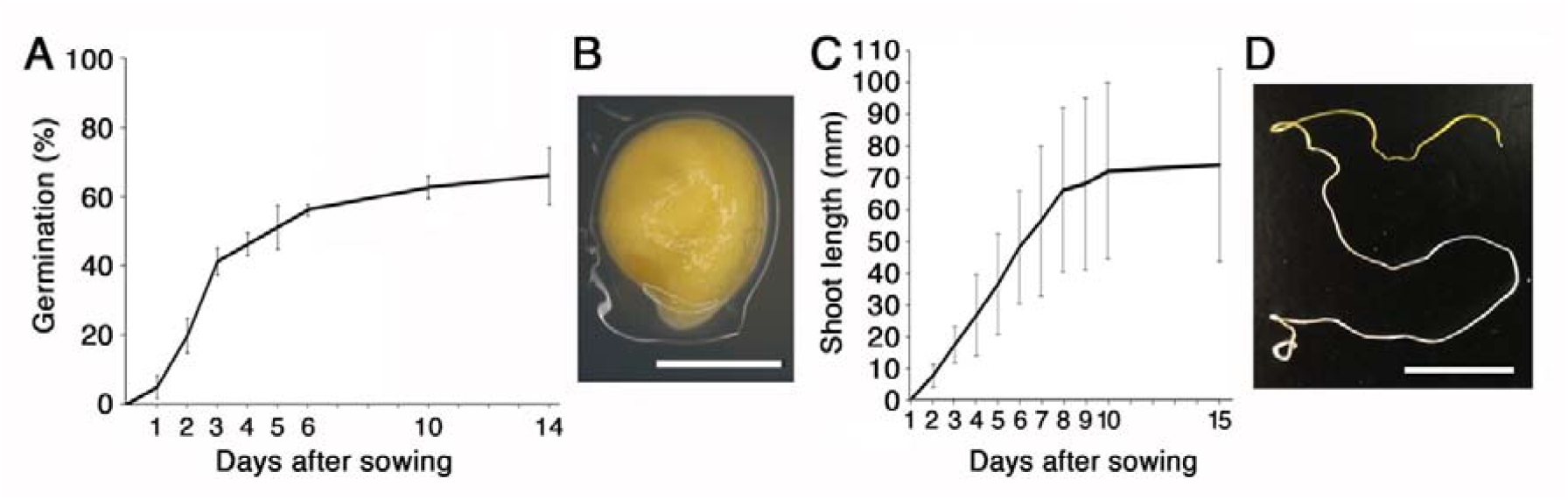
The rate of germination and shoot growth of *C. campestris*. **A** Time course of the germination rate of the seeds, which were sown on MS agar plates and placed under 16/8 h white light/dark cycle at 25 °C. Data are means ± SE (n = 3, 20 seeds each). **B** Morphology of the seed that begins root protrusion and is judged germinated. Bar = 1 mm. **C** Time course of the shoot length after the seed sowing. Data are means ± SE (n = 10). **D** Whole seedling morphology at 10 days after sowing. Bar = 1 cm.

We first examined the effect of light conditions on seed germination and shoot growth. When the plates were placed under continuous white, red, far-red, blue light or darkness for 3 days after sowing, germination was enhanced by far-red, blue, and no light conditions but rather reduced by red light in comparison with that under white light (Fig. 2A). We then examined the effect of alternate incubation with red and far-red light for 24 h each on germination and found that the latter illumination is effective (Fig. 2B), suggesting the involvement of reversible phytochrome regulation. After transfer of 3-day-old seedlings germinated under white light, shoot growth was examined under different light conditions. While the haustorium formation was confirmed to be induced by far-red light (Fig. 2C), the shoot length was rather shortened under far-red light compared with other light conditions (Fig. 2D)

**Fig. 2.**
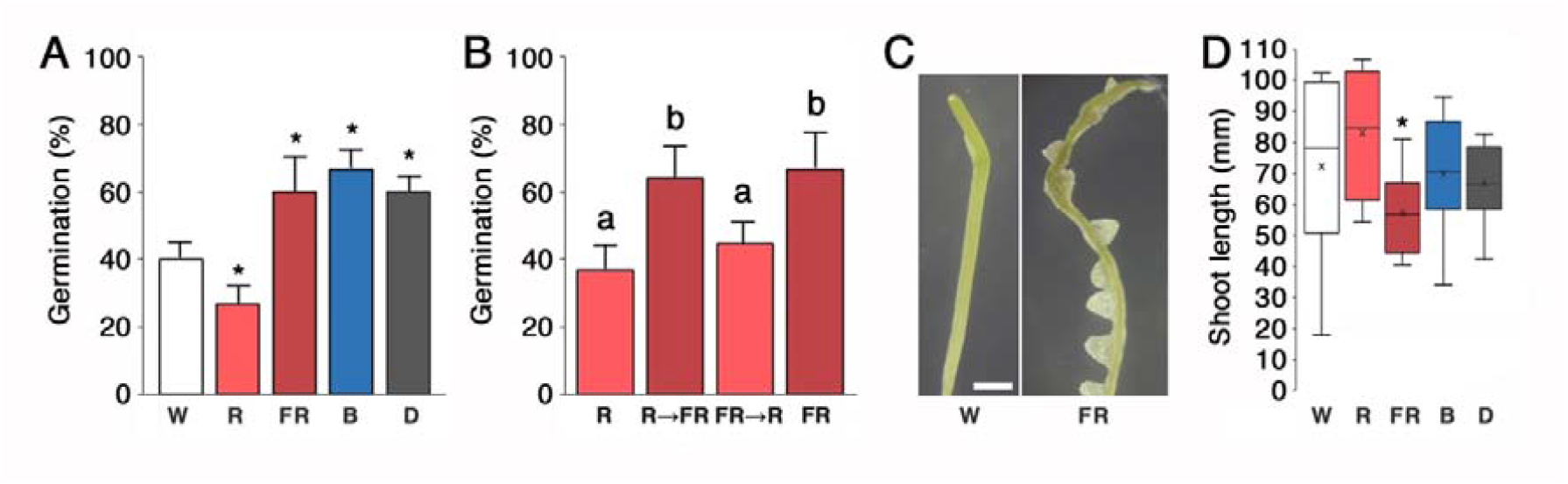
Effect of different light conditions on germination and shoot growth. **A** Effect of light wavelengths on germination. The germination rate was measured at 3 days after the seeds were sown on MS plates and placed under continuous white (W), red (R), far-red (FR), blue (B) light or darkness (D). Data are means ± SE (n = 3, 20 seeds each). Asterisks indicate a significant difference from the white-light treatment (*p* < 0.05). **B** Effect of red and far-red light treatments on germination. The imbibed seeds were placed under red light for 48 h (R), red light for 24 h followed by far-red light for 24 h (R→FR), far-red light for 24 h followed by red light for 24 h (FR→R), or far-red light for 48 h (FR). Each treatment was followed by the dark treatment for 24 h. Data are means ± SE (n = 3, 20 seeds each). Different letters indicate a significant difference (*p* < 0.05) between treatments. **C** Shoot morphology of 10-day-old seedlings grown under white (W) and far-red (FR) light. Bar = 5 mm. **D** Effect of light wavelengths on shoot elongation. Three-day-old seedlings grown under 16/8 h white light/dark were transferred under continuous white (W), red (R), far-red (FR), blue (B) light or darkness (D) and grown for 7 days. Data are means ± SE (n = 10). An asterisk indicates a significant difference from the white-light treatment (*p* < 0.05).

### Alkaline pH limits seed germination

When the seeds were placed on agar media with different pH conditions, germination was remarkably reduced at pH higher than 7.0 (Fig. 3A). As for temperature, higher temperature below 30 °C was more effective on germination (Fig. 3C). We also examined the effect of sugars and high salts on germination. Supplementation of 100 mM sucrose, glucose, fructose, maltose, or mannose had no significant effect on both the germination efficiency and the shoot elongation (Fig. 3E, F). NaCl and KCl had an inhibitory effect on germination significantly at 50 mM and at 200 mM, respectively (Fig. 3G). On the other hand, exposure of the seeds to mannitol, used as an osmotic control, had an inhibitory effect on germination at 500 mM while water submergence completely inhibited the seed germination (Fig. 3G).

**Fig. 3.**
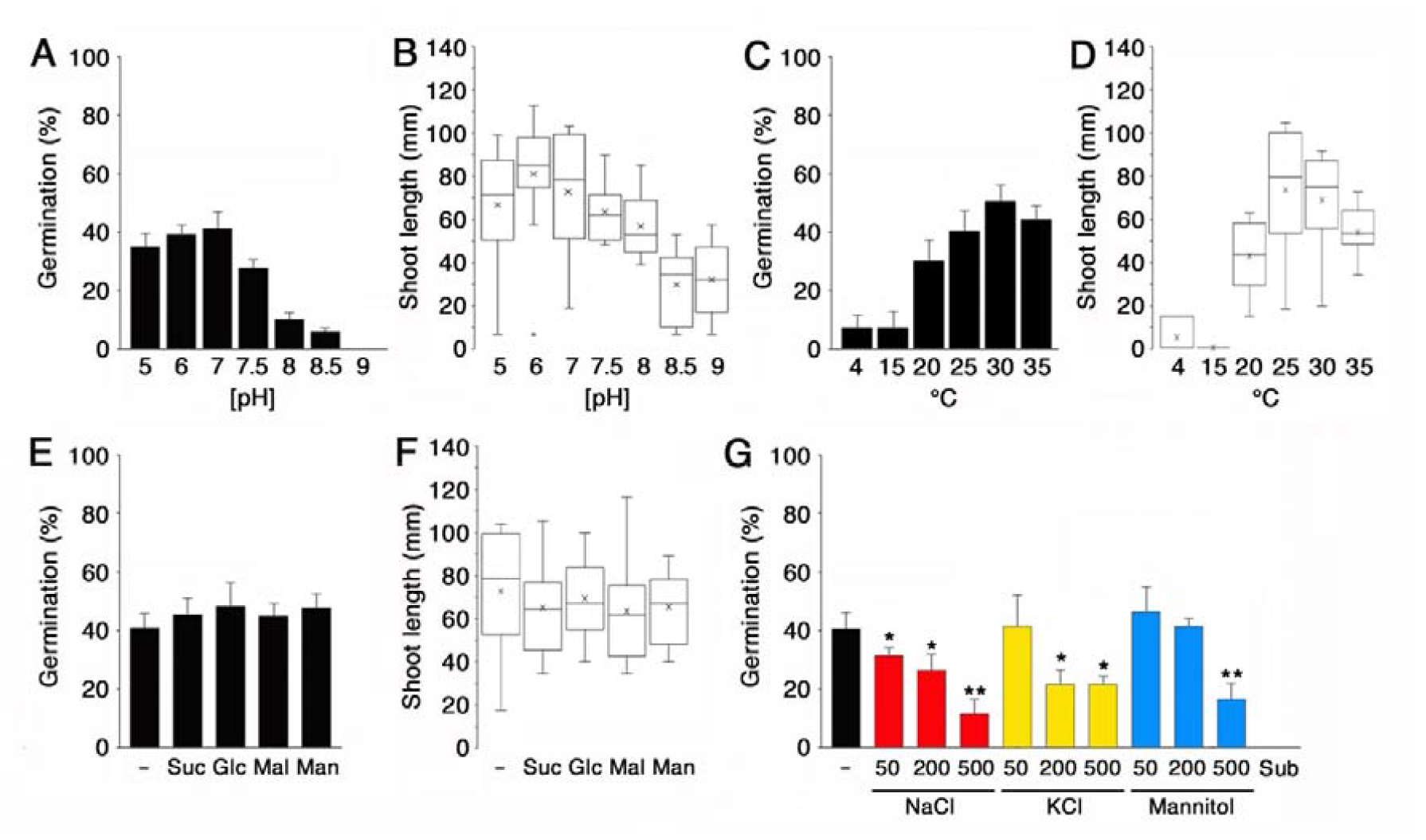
Effect of pH, temperature, sugar, and high salt conditions on germination and shoot growth. **A** and **B** Effect of media pH on germination (**A**) and shoot elongation (**B**). **C** and **D** Effect of temperature on germination (**C**) and shoot elongation (**D**). MS agar plates were upside down. **E** and **F** Effect of sugars on germination (**E**) and shoot elongation (**F**). 100 mM sucrose (Suc), glucose (Glc), maltose (Mal), or mannose (Man) was added to MS agar plates. **G** Effect of high salt and osmotic stresses on germination. Data are means ± SE (n = 3, 20 seeds each in **A, C, E**, and **G**, n = 10 in **B, D**, and **F**). In **B, D**, and **F**, 3-day-old seedlings grown in mock conditions were transferred to the indicated conditions and grown for 7 days. In **G**, asterisks indicate a significant difference from the mock (-) treatment (* *p* < 0.05, ** *p* < 0.01). Sub, submergence by water.

### Effect of phytohormones on germination and growth

Among the hormones examined, supplementation of 1 µM gibberellic acid (GA3) enhanced germination and that of 5 µM abscisic acid (ABA) reduced it (Fig. 4A). In addition, methyl jasmonate (MeJA) was shown to enhance germination at 1 µM (Fig. 4A). When the seedlings were transferred from MS plates to those with different hormones after germination, the shoot elongation was significantly enhanced by MeJA but repressed by 5 µM indole acetic acid (IAA), ABA, 100 µM amino-cyclopropane carboxylic acid (ACC), a precursor of ethylene, and 100 µM salicylic acid (SA) (Fig. 4B). Bikinin, an inhibitor of GSK3 kinases, which is used as a substitute for brassinosteroids (De Rybel et al. 2009), solely had no obvious effect on germination and shoot growth.

**Fig. 4.**
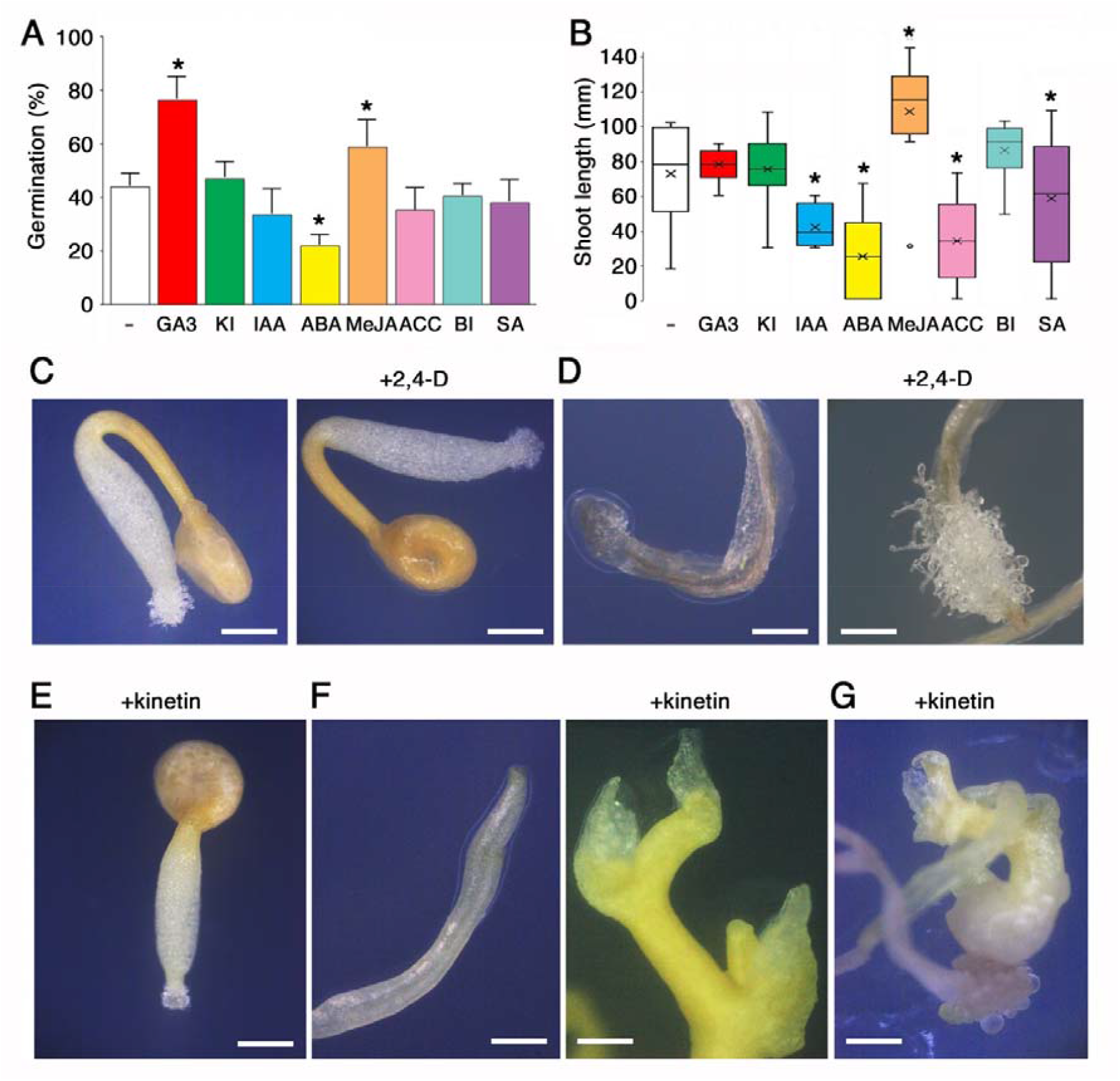
Effect of phytohormones on germination and shoot growth. **A** and **B** Effect of phytohormone treatments on germination (**A**) and shoot elongation (**B**). Phytohormones and their substitutes examined were 1 µM gibberellic acid (GA3), 1 µM kinetin (KI), 5 µM indole acetic acid (IAA), 5 µM abscisic acid (ABA) 1 µM methyl jasmonate (MeJA), 100 µM amino-cyclopropane carboxylic acid (ACC), 100 µM bikinin (BI), and 100 µM salicylic acid (SA). In **B**, 3-day-old seedlings grown in mock conditions were transferred to the indicated conditions and grown for 7 days. Data are means ± SE (n = 3, 20 seeds each in **A**, n = 10 in **B**). Asterisks indicate a significant difference from the mock (-) treatment (*p* < 0.05). **C** Morphology of 4-day-old seedlings grown without or with 5 µM 2,4-D. **D** Morphology of the root of 10-day-old seedlings grown without or with 5 µM 2,4-D. **E** Morphology of 4-day-old seedlings grown with 1 µM kinetin. **F** Morphology of the shoot of 10-day-old seedlings grown without or with 1 µM kinetin. **G** Morphology of the shoot of 20-day-old seedlings grown with 1 µM kinetin. Bars in **C** to **G** = 1 mm.

We also examined the effect of an artificial auxin, 2,4-D on the seedling growth and found that 5 µM 2,4-D prolonged the survival of trichomes at the root tip, which are normally decayed in 10 days after germination (Fig. 4C, D). Treatment with 1 µM kinetin caused no seedling hook formation (Fig. 4C, E). It is also noted that kinetin treatment induced formation of scale leaves on the shoot and the emergence of lateral shoots in 10-day-old seedlings (Fig. 4F). Its treatment for 20 days resulted in the callus formation at the base of the shoot and the root tip (Fig. 4G).

### Responses of seeds and seedlings to amino acids and polyamines

We next investigated the effect of exogenous amino acids on germination and shoot growth. Among 20 amino acids, alanine and aspartate slightly but significantly increased the germination rate and histidine reduced it (Fig. 5A). The germination rate at 10 days after sowing was not different between the seeds treated with mock, alanine, and aspartate, but that in the presence of histidine remained low (Fig. 5B). All the amino acids examined had no significant effect on the shoot elongation (Fig. 5C).

**Fig. 5.**
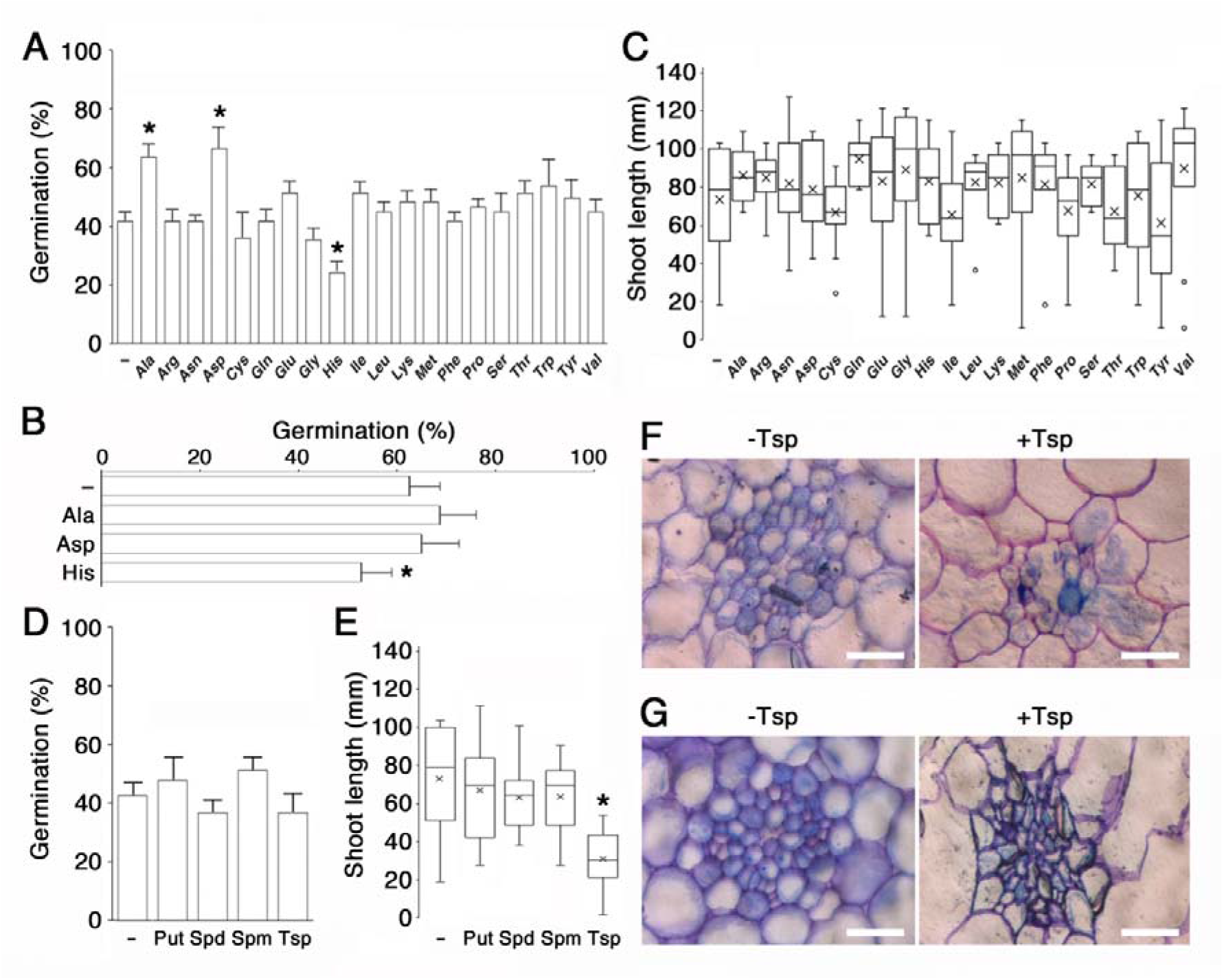
Effect of amino acids and polyamines on germination and shoot growth. **A** and **B** Effect of amino-acid treatments on germination. The germination rate was measured at 3 days (**A**) and 10 days (**B**) after the seeds were sown on MS plates with 1 mM of the indicated amino acid. Data are means ± SE (n = 3, 20 seeds each). Asterisks indicate a significant difference from the mock treatment (*p* < 0.05). **C** Effect of amino-acid treatments on shoot elongation. Three-day-old seedlings grown in mock conditions were transferred to the indicated conditions and grown for 7 days. Data are means ± SE (n = 10). **D** and **E** Effect of polyamine treatments on germination (**D**) and shoot elongation (**E**). 1 mM putrescine (Put), 1 mM spermidine (Spd), 100 µM spermine (Spm), or 100 µM thermospermine (Tsp) was added to MS agar plates. In **E**, 3-day-old seedlings grown in mock conditions were transferred to the indicated conditions and grown for 7 days. Data are means ± SE (n = 3, 20 seeds each in **D**, n = 10 in **E**). An Asterisk indicates a significant difference from the mock (-) treatment (*p* < 0.05). **F** and **G** Sections of the root (**F**) and the basal part of the shoot (**G**) of 10-day-old seedlings grown without or with 100 µM thermospermine. Bars = 100 µm.

We further found that treatment with polyamines caused no apparent effect on the germination (Fig. 5D) while thermospermine had an inhibitory effect on the shoot elongation (Fig. 5E). A structural isomer of spermine, thermospermine, plays a role in negatively regulating xylem differentiation (Takano et al. 2010). To examine the effect of thermospermine on the vascular development, we observed sections of the seedlings and found that differentiation of vascular cells was severely repressed by thermospermine in the basal part of the shoot (Fig, 5F). Sections of the root revealed that, although the development of vascular tissues were not affected by thermospermine, the shape of cells was overall getting distorted in the seedling treated with thermospermine (Fig, 5G), suggesting an accelerated degeneration of the root tissue.

### Environmental responses of gene expression in the seedling

We finally addressed whether the molecular responses to external stresses have been evolutionarily retained in *C. campestris* seedlings or not. qRT-PCR experiments revealed that expression of an *RbcS* homolog, which putatively encodes a small subunit of Rubisco in the chloroplast, was not up-regulated after transfer from dark to white light while a homolog of *Phytochrome A* (*PhyA*), whose expression is generally up-regulated in the dark, showed an increased expression by 24-h dark treatment (Fig. 6). We confirmed that expressions of homologs of an ABA-responsive *SPMS* encoding spermine synthase (Yariuchi et al. 2021), a salicylic acid (SA)-responsive *VSP2* (Koornneef et al. 2008), an MeJA-responsive *WRKY18* (Potschin et al. 2014), and a heat-shock protein gene, *HSP90*, were respectively increased by treatment with ABA for 24 h, SA for 24 h, MeJA for 24 h, and heat stress at 37 °C for 4 h (Fig. 6).

**Fig. 6.**
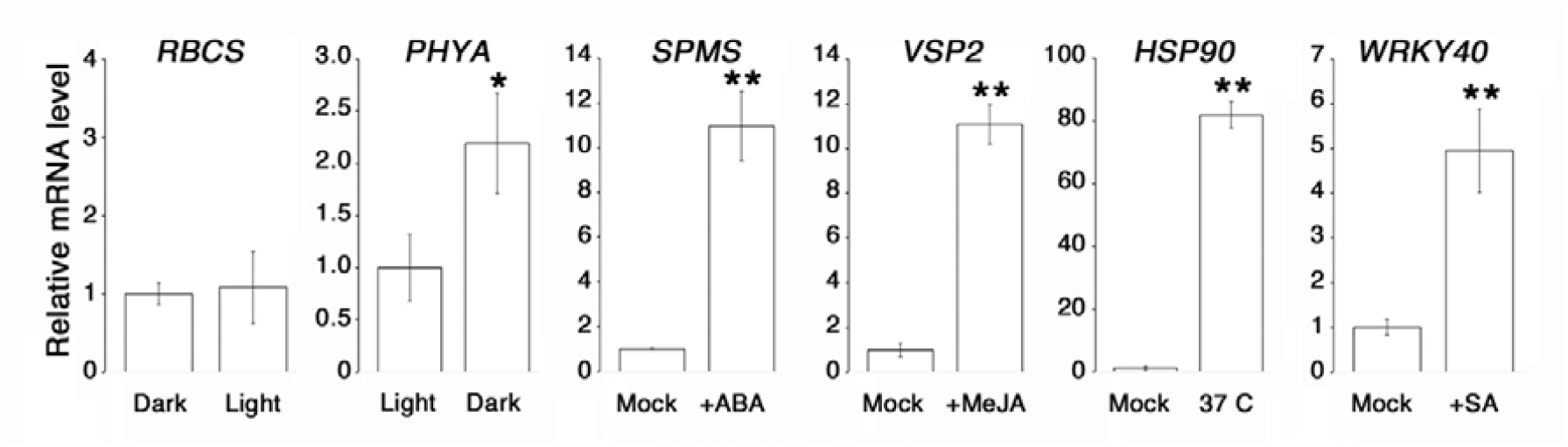
Effect of light, high temperature, ABA, MeJA, and SA on the expression of the genes known to be responsive in other plants. Expressions of putative *C. campestris* homologs of *RbcS* (accession #VFQ99940), *PhyA* (accession #VFQ60648), *Spms* (accession #VFQ58264), *Vsp2* (accession #VFQ80521), *Hsp90* (accession #VFQ72691), and *Wrky40* (accession #VFQ87276) were examined by qRT-PCR. Light treatments were 24-h white light on 9-day-old dark-grown seedlings in MS agar plates and vice versa. Other treatments were 3-h 5 µM ABA, 12-h 1 µM MeJA, 3-h heat shock at 37 °C, and 12-h 100 µM SA on seedlings grown in MS plates for 9 days and preincubated for 24 h in MS liquid media. *EF1a* (accession #VFQ91475) was used as the reference gene. Transcript levels relative to the mock treatment are shown. Data are means ± SE of three different experiments. Asterisks indicate a significant difference from the mock treatment (* *p* < 0.05, ** *p* < 0.01).

## Discussion

Light quality and duration are pivotal in triggering the twining response and haustoria formation in dodders (Lane and Kasperbauer 1965; Furuhashi et al. 1995; Tada et al. 1996; Orr et al. 1996; Haidar 2003). A study in *C. japonica* has revealed that blue light is essential for twining and a lower red/far-red light (R/FR) ratio is important for subsequent haustoria induction (Furuhashi et al. 2021). Blue light-dependent twining may be a variant of blue light-mediated phototropic responses. Both far-red light and touch stimuli play a critical role for the haustoria formation as a sign showing the presence of putative host plants. Low R/FR irradiation has been shown to induce the expression of genes related to cell wall degradation in *C. chinensis* (Pan et al. 2022). Stage-specific genes during far-red light-induced haustoriogenesis have been identified in *C. campestris* (Bawin et al. 2022). Another study in *C. campestris* has shown that mature shoots exhibit coiling around the host and haustoria formation even under red light, suggesting that the response to red light is modified during the transition from young to mature shoots to facilitate parasitism (Yokoyama et al. 2023). In the present study, we found that seed germination is also stimulated by far-red light and reduced by red light. We further confirmed that, although not completely, the germination response to red and far-red light is reversible, suggesting the involvement of phytochrome signaling in germination. This response is apparently opposite to that observed in many plant species whose germination is induced or promoted by a single red-light pulse. Germination in some plant species requires only rehydration in the dark during which phytochromes may be converted into the active far red-absorbing form (Pfr). In tomato, germination is inhibited not only by far-red light but also by strong continuous red light (Shichijo et al. 2001). This inhibition is suggested to be caused by degradation of the active but light unstable Pfr of PhyA (Appenroth et al. 2006). Given the reversed response to red and far-red light, *C. campestris* seeds may possess a unique phytochrome signaling mechanism for germination against the conventional pathway of both light and dark germinators. Far red-light induced germination was first reported for a bromegrass, *Bromus sterilis* (Hilton 1982). Interestingly, the similar response is observed in another holoparasitic plant, *Orobanche coerulescens* (Takagi et al. 2009). Thus, the mechanism may have been evolved independently in some families, although the causal relationship between the evolution of parasitism or the loss of photosynthesis and the reversal of the phytochrome action remains unknown. Shoot elongation is also reduced by far-red light in *O. coerulescens* (Takagi et al. 2009), suggesting a common alteration of phytochrome signaling components in these parasites. Alternatively, the reduction in the shoot length under far-red light observed in *C. campestris* might be explained by a trade-off with the haustoria formation.

Among abiotic stresses examined in this study, the pH higher than 7 was found to have a severely inhibiting impact on germination. A similar response to alkaline conditions and the preference to acidic conditions in seed germination have been reported for some species of the *Ipomoea* genus (Oliveira and Norsworthy 2006; Singh et al. 2012). Hence, the alkaline soil pH is likely to be a limiting factor for germination generally common to the morning glory family, Convolvulaceae. Our results showing that the germination rate reached the highest at 30 °C and was markedly reduced below 20 °C are consistent with previous reports on the effect of temperature on filed dodder germination (Hutchinson and Ashton 1980; Benvenuti et al. 2005; Goldwasser et al. 2016). A similar tendency was observed in the shoot elongation. These traits seem to reflect a tropical origin of dodders.

It is noteworthy that, in hormone experiments, MeJA promoted both seed germination and shoot elongation. Many studies have revealed that jasmonates induce defense responses against pathogens but have negative effects on plant growth and germination (Linkies and Leubner-Metzger 2012; Yang et al. 2012; Ghorbel et al. 2021; Sohn et al. 2022; Pan et al. 2023). The level of jasmonates is closely correlated with the accumulation of ABA in *Arabidopsis* seeds (Dave et. al. 2016). JASMONATE RESISTANT 1 (JAR1), which encodes an enzyme to produce a bioactive molecule, jasmonoyl-isoleucine in *Arabidopsis*, and is also named FAR-RED INSENSITIVE 219 (FIN219), participates in PhyA-mediated far-red light signaling (Chen et al. 2015). It is thus possible that JAR1/FIN219 plays a role in germination activated by far-red light and MeJA in *Cuscuta*. There are, however, some plants that show jasmonate-stimulated germination, such as pear (Yildiz et al. 2008), apple (Górnik et al. 2018) and wheat (Xu et al. 2016). Further research will be necessary to evaluate the mechanism of jasmonate signaling and its crosstalk with light and other hormone pathways in *Cuscuta*.

Our results also revealed that *C. campestris* can develop scale leaves without haustoria formation by treatment with kinetin. The degree of development of scale leaves may be different among *Cuscuta* species. The shoot tip of *C. japonica, C. australis*, and *C. chinensis* has several spirally arranged minute scales (Fujita 1964). Although a study indicated that the seeds of *C. campestris* contains approximately 1 µM zeatin (Al-Gburi et al. 2019), it remains to be examined whether the degree of scale leaf development is attributed to the endogenous level of cytokinin or not. Taking previous research into account (van der Kooij et al. 2000; Vogel et al. 2018), retrogression of the leaf development may be tightly associated with the level of host dependence or the defective level of photosynthesis.

We further found that thermospermine drastically repressed the development of xylem vessels in elongating shoots, confirming the function of thermospermine in negatively regulating xylem formation (Takano et al. 2012). However, we observed no such effect in the root section but instead the distortion of cell shapes. This is probably because the root part formed during embryogenesis shows little or no division and differentiation of cells after germination in this plant. Distorted cell shapes represent a symptom of tissue degeneration. Reduced xylem development in the shoot could cause reduced absorption of water and nutrients, and consequently accelerate the root tissue degeneration and also inhibit the shoot elongation.

On the other hand, among amino acids examined, alanine and aspartate accelerated germination. Alanine is converted with 2-oxoglutarate to pyruvate and glutamate by alanine aminotransferase (AlaAT/ALT). The long seed dormancy in wild-type barley has been shown to be linked to the recessive allele of *AlaAT* (Sato et al. 2016). A positive effect of alanine on germination detected in *Cuscuta* might be due to the function of AlaAT in breaking dormancy. Aspartate is converted with 2-oxoglutarate to oxaloacetate and glutamate or is conversely synthesized by Aspartate aminotransferase (GOT1/AAT). This enzyme plays a key role in regulating carbon and nitrogen metabolism in most organisms. During *Arabidopsis* germination, aspartate is increased the most among all kinds of amino acids (Fait et al. 2006). Given that both AlaAT and GOT1 are directly involved in providing substrates for TCA cycle (Han and Yang 2015), it is possible that supplementation of alanine and aspartate is effective to generate sufficient energy via the respiratory chain for germination. Or otherwise, pyruvate produced by AlaAT and aspartate could serve as a substrate for gluconeogenesis (Walker et al. 2021) and sugar supply via the glyoxylate cycle (Graham 2008), respectively. In contrast, only histidine was shown to be inhibitory to germination.Histidine promotes plant seed oil deposition through ABA biosynthesis and β-oxidation (Ma and Wang 2016). It remains to be determined whether exogenous histidine activates genes of ABA biosynthesis in imbibed seeds of *Cuscuta* or not. Considering a role of histidine in metal ion chelation (Seregin and Kozhevnikova 2021), metal ion homeostasis might be seriously affected by excess histidine.

In conclusion, the results of our study revealed unique responses to environmental conditions in *C. campestris* seeds and seedlings before parasitism. Further experimental work will increase our insight into the whole lifestyle and the (dis)advantage of parasitic strategy of dodders.

## Acknowledgements

KN thanks the Public Interest Incorporated Foundation ”Ohmoto Ikueikai” for the generous financial support.

## Author Contributions

KN conducted all experiments, compiled the data with statistical analyses, and wrote the draft; TT and RY conceptualized, planned, and supervised the experiments, and undertook critical editing of the manuscript. All the authors approved the final version submitted.

## Data Availability

All the data are provided as Figures and Online Resource (Supplementary Figure).

## Declarations

The authors have no relevant financial or non-financial interests to disclose.

