## Supplementary Fig for "Effect of environmental conditions on seed germination and seedling growth in *Cuscuta campestris*"

**Supplementary Fig. 1** Primer sequences used in qRT-PCR experiments and their surrounding genomic DNA sequences. Primer sequences are in bold and protein coding sequences are framed. Long intron sequences are omitted by asterisks.

EF1A (accession #VFQ91475)

EF1A-F: **CTGCTGCAACAAGATGGATG**

EF1A-R: **TCTTCTTCTGGGCAGCTTTG**

CAAATGATTTG**CTGCTGCAACAAG**GTTTGTTGTTATGACCCACTGTTTGTTTTCAGTCTTAGTCTTGATGTATTTAATATGAACCTAGTTTGTTTACCCAAGTATTTGATATGAACCTTGTTTGTTTCCCTTTGTGATGTCTGACAG**ATGGATG**CCACTACCCCCAAATACTCCAAGGCCAGATATGATGAAATTGTGAAGGAGGTGTCTTCATACCTGAAGAAGGTTGGGTACAACCCTGATAAAGTTCCCTTTGTCCCCATCTCTGGATTTGAGGGAGACAACATGATTGAGAGGTCAACCAACCTTGACTGGTACAAGGGCCCAACACTTCTGGAAGCTCTTGATCAGATCAATGAGCCAAAGAGGCCTTCTGACAAGCCACTCCGTCTCCCACTTCAGGATGTTTACAAGATTGGTGGTATTGGAACTGTCCCTGTTGGACGTGTTGAGACTGGTGTTATCAAGCCTGGTATGGTTGTGACCTTTGCCCCTTCTGGCTTGACAACTGAAGTTAAGTCTGTTGAGATGCACCACGAGGCTCTGACTGAAGCACTCCCTGGAGACAATGTTGGGTTTAACGTGAAGAATGTTGCTGTGAAGGATCTGAAGCGTGGGTATGTTGCCTCCAACTCCAAAGATGATCCTGCCAAGGAAGCTGCCAATTTCACCTCCCAGGTCATCATCATGAACCATCCTGGTCAGATTGGAAATGGATATGCCCCTGTGCTCGATTGCCACACATCCCACATTGCTGTCAAGTTTGCTGAGCTTGTCACCAAGATTGACCGAAGGTCTGGCAAGGAGCTCGAGAAGGAGCCGAAGTTCCTCAAAAATGGTGACGCTGGTTTTGTTAAGATGATTCCCACCAAGCCAATGGTTGTCGAGACCTTCGCTGAGTACCCGCCGCTTGGTCGGTTTGCCGTGAGGGACATGCGTCAAACAGTTGCTGTTGGTGTCATCAAAGCTGTGGAGAAGAAGGACCCGACAGGAGCAAAAGTCAC**CAAAGCTGCCCAGAAGAAGA**AATGAGCAATTGTTTCCGACCTCTAG

RBCS (accession #VFQ99940)

RBCS-F: **TTCCCCATCTCCAAGAAGTC**

RBCS-R: **CGAATGGATTCTCCAACTCG**

GCCGCC**TTCCCCATCTCCAAGAAGTC**CGCCGCCGACACCGCTTCCTTGGCCACCAACGGTCTCAGAGTACAGTGTATGAAGGTATCTAATTAGGCTATTTCATCCACTTATGTGTTTCATT*****

*****ATGTACAAATGAATACAAACAGGTGTGGCCACCGGCGGAAAACAAGAAGTTTGAGACATTGTCATACCTCCCAGACTTGACCGACGAGCAGTTGTATAAACAAGTGGAGTACCTCCTCAGGAATCAATGGATCCCTTGCTTGGAATT**CGAGTTGGAG**GTATAGGTCATGTCTCGTATACTCGTACGCGTT*****

*****ACATTTGATGGAATATATATGTCTGTCATGTAG**AATCCATTCG**TGCAACGCGAAAACC

PHYA (accession #VFQ60648)

PHYA-F: **CTTGATCAAGTGTTGGTTGC**

PHYA-R: **CTGTGTCTCACTCGTAATTC**

AGTTCAAA**CTTGATCAAGTGTTGGTTGC**TTCAATCAGTCAAGTGATGGCAAAGAGCAATGGGAAATGTTTGAGGATAACTAATGACATGTGTGAGAGTGTTGTACATGAAACATTGTATGGAGACTCTCTAAGGCTTCAACAAATCCTTGCCGAATTTTTGTCAGTAGCCGTGGATTACACCCCGAGGGGAGGCCAGCTTGATCTTTCATCTGCATTAACCAAAGATCTTTTGGGAGAATCCGTTCGTCTTGCCCGTTTG**GAATTACG**GTACGATAT*****

*****AATTTTACTTTTTAAAAAATGTGTTGTTGAAATTCAG**AGTGAGACACAG**CGGGAGGGGGG

SPMS (accession #VFQ58264)

SPMS-F: **GGTGTTCTTTGCAACATGGC**

SPMS-R: **CTCTGTGAACCTCAGAATTG**

GAGACATTAGCTAGGGCATTAAGACCCGGT**GGTGTTCTTTGCAACATGGC**AGAGAGCATGTGGCTTCACACACATTTGATTCAGGATGTAATTTCAATTTGCCGCGAAACATTCAAGGGCTCTGTCCGGTATGCATGGACCAGCGTCCCTACGTATCCAAGGTAATAGTTCCCAAAACCAGTCAAGTGACTTAGGTTTC*****

*****GAGGAACTTGATTTTTAGTTGTCAAGGATAGATAATCATGCATATTGTGTGCAGTGGTGTTATCGGGTTCTTGTTATGCTCAACTGAAGGCCCACCTGTTGATTTTTTGAACCCAATAAATCCCATTGAGAAGCTTGAGGGAGCCCTCCAATATCGCCGCCAACTAAGGTTTTA**CAATTCTGAG**GTATGTATGCTAT*****

*****CATTTATGTTGGGATCGACAG**GTTCACAGAG**CGGCATTTGCCTTGCCGGC

VSP2 (accession #VFQ80521)

VSP2-F: **CAACATGGTTTTGGGTTGGAG**

VSP2-R: **ACGGCTGCCGATTTTACATG**

TGGACGAGACTTTGATTTCCAATCTCCCTTATTACTCT**CAACATGGTTTTGG**GTAAGTTTTA*****

*****TTTGATTAAATGTTGTTGGGTGATTGTGAGTGATTGCAG**GTTGGAG**GTTTTTGATGGGGTGGAGTTTGACAAGTGGGTTGAGAAGGGAAGAGGGCCTGCGATAAACTCAAGCTTGAAGCTTTATCAAGAAATCAAAGAATTGGGATTCAAAGCTTTCTTGTTGACAGGAAGGAGTGATAGACACAGAGAAGTCACTGAGGAGAATTTGGTAAATGCTGGTTTTCAAGATTGGGATAAATTCATTCTTAGGTGAGTCTCATTCCACTATTTGA*****

*****TTTCAATTTTTTCTTACAGGTCAGCGGAGGAC**CATGTAAAATCGGCAGCCGT**GTACAAATCGGAGA

HSP90 (accession #VFQ72691)

HSP90-F: **AGGACCAGTTGGAGTACTTG**

HSP90-R: **GCCATTCATGGAGAGACCTC**

GATGTAGATGGAGAACCGCTTGGAAGGGGAACCAAGATCACTCTCTTCCTCAAGG**AGGACCAG**GTAAATAATTTTGCAATTCAATAATTTCTGAATTATAATGCGTGTTAATGATTGTGCAAGCATCTTGACTTATGTTTGATATTTTGCAG**TTGGAGTACTTG**GAGGAGAGGCGACTCAAGGACCTCGTGAAGAAGCATTCTGAGTTCATTAGCTACCCAATCCACCTGTGGACCGAGAAGACAACTGAGAAGGAAATCAGTGATGATGAGGATGACGAGCCCAAGAAGGAAGAAGAAGGCGATATTGAGGAAGTTGATGAGGAAAAGGAGAAAGAAGGTAAAAAGAAGAAGAAGATCAAG**GAGGTCTCTCATGAATGGC**AGCTCGTCAACAAACAAAAACCACTCTGGCTGCGGAAGCCAGAAG

WRKY40 (accession #VFQ87276)

WRKY40-F: **GTCAAGAAAAAGGTCCAAAGAAG**

WRKY40-R: **GAATGGATTTGGCGATGCCAG**

TGCTCTTTTGCCCCTACTTGCCCT**GTCAAGAAAAAG**GTAAAATTTCTCTTTTGTAGTT*****

*****ATTTGTTTGAATTTTAAACATATTTCAG**GTCCAAAGAAG**CATTGAAGATCAGTCCATTTTGGTGGCCACCTACGAAGGGGAACACAACCACCCCTCAAAGCCGGACCAACCCGCCGCCGGCGCTTCCACGGCGGCCAACCGTTCCATCGCGACGCCCCGCGGTGGATCCGCCGCCTCCACAATCACCAGCCCCGGAAAACCAGCCACCCCCCTGGAT**CTGGCATCGCCAAATCCATTC**CCGCCCGGCAGAGTCA
